# A simple, consistent estimator of heritability for genome-wide association studies

**DOI:** 10.1101/204446

**Authors:** Armin Schwartzman, Andrew J. Schork, Rong Zablocki, Wesley K. Thompson

## Abstract

Analysis of genome-wide association studies (GWAS) is characterized by a large number of univariate regressions where an outcome, a quantitative trait, is regressed on hundreds of thousands to millions of genomic markers, i.e. single-nucleotide polymorphism (SNP) counts, one marker at a time. Assuming a linear model linking the markers to the outcome, this article proposes an estimator of the heritability of the trait, defined here as the fraction of the variance of the trait explained by the genomic markers in the study. The estimator, called GWAS heritability (GWASH) estimator, is easy to compute, highly interpretable, and is consistent as the number of markers and the sample size increase. More importantly, it can be computed from summary statistics typically reported in GWAS, not requiring access to the original data. The estimator takes full account of the linkage disequilibrium (LD) or correlation between the SNPs in the study through moments of the LD matrix, estimable from auxiliary datasets. Unlike other proposed estimators in the literature, the precision of the estimate is obtainable analytically, allowing for power and sample size calculations for heritability estimates.

## 1 Introduction

Modern statistics is increasingly seeing situations where there is an interest in understanding how an outcome measured on a large number of subjects is related to a much larger number of predictors. Genome-wide association studies (GWAS) are an excellent example of this, where the outcome is an observed trait or phenotype and the predictors are single-nucleotide polymorphisms (SNPs), each captured by the number of copies of the reference allele (0, 1 or 2). The sample size in typical GWAS may be in the order of tens to hundreds of thousands, while the number of SNPs may be ten to one hundred times as large, in the order of millions.

Suppose the outcome is related to the predictors by a linear model. Since the number of coefficients in the model is larger than the sample size, this is an under-determined problem and it is impossible to estimate the coefficients without additional assumptions. Low dimensional summaries, however, are estimable. In particular, in this paper we focus on the heritability, defined as the proportion of variance of the outcome explained by the measured predictors. In the GWAS literature this is often referred to as chip heritability or SNP heritability [24]. Heritability is an important parameter in that it helps quantify the proportion of the observed outcome that can be predicted from genetic factors. In other words, the estimation of heritability is often the first step in investigating the genetic basis of traits and diseases.

A prototypical GWAS aims to identify individually important genetic factors by regressing an outcome variable on each SNP, one at a time, selecting only the most stringently significant SNPs (*p* < 5 × 10^−8^) as discoveries. The basic GWAS approach has become a mainstay in the genetic analysis of complex traits. Thousands of studies have been performed and tens of thousands of candidate causal variants have been cataloged for all variety of trait and disease (www.ebi.ac.uk/gwas/, [16]). In part due to funding institution data sharing mandates, to increase transparency and to fuel post-hoc and secondary analysis of GWAS results [18], per-SNP univariate regression statistics (beta coefficients, *t*-statistics, *p*-values, standard errors, etc.) are now regularly published along with GWAS articles. While privacy concerns often prevent the sharing of subject level genotypes and phenotypes, these summary statistics are readily available for hundreds of individual GWAS studies (e.g., www.ebi.ac.uk/gwas/downloads/summary-statistics).

It is of interest, therefore, to develop an estimator of heritability that can provide accurate estimates using only summary statistics from GWAS. Computational efficiency is another desired property given the large size of the data. And, as with any estimation procedure, interpretability is also desired in order to gain further insights into the data. For example, we wish to understand how heritability is affected by the correlation between the predictor genomic markers, called linkage disequilibrium (LD) in the context of GWAS.

In this paper, we propose an estimator called GWAS heritability (GWASH) estimator The estimator is based on the variance-fraction estimator in [6] and is astonishingly simple. For a GWAS with *m* predictors and *n* independent subjects, the estimator is

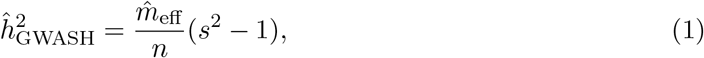

where *s*^2^ is the empirical variance of the GWAS t-statistics (up to a small transformation) and 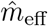 is the estimated effective number of SNP predictors. The latter is equal to the number of predictors *m* divided by a factor 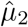, an estimate of the second spectral moment of the LD matrix that captures in a single number the effect of LD: the higher the LD, i.e. the higher the correlation between the predictors, the lower their effective number.

The GWASH estimator is not only easy to remember and compute as a simple formula. It also has an interpretation as being proportional to excess empirical variance of the univariate t-statistics with respect to the complete null hypothesis of independence between the outcome and the predictors, in which case the the empirical variance is about 1. The empirical variance *s*^2^ is in itself an intuitive quantity that summarizes the strength of the relationship between the predictors and the outcome, and has been used as a simple measure of signal to noise ratio in GWAS contexts [19]. Thus, the proposed estimator has the nice property that it increases linearly with enrichment, where the proportionality constant depends on LD.

Moreover, the formula dictates that LD affects the estimation of heritability as a scaling factor, yielding a definition of the effective number of SNPs involved. Computing the factor 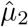 to find the effective number of predictors is the only relatively difficult part of the estimation. The factor 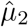 estimates 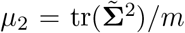, where Σ̃ is the correlation matrix of the predictors, the LD matrix. As a first approximation, the patterns of correlations among SNPs can be taken as a feature of a given population and estimated from publicly available data resources such as the 1000 genomes project (1KGP) [10] (http://www.internationalgenome.org/). This approach has been reasonable when the reference sample plausibly represents the same population as the GWAS sample in contexts including imputation [13], heritability estimation (e.g., [2]), functional fine-mapping (e.g., [21]) and various post-hoc burden tests (e.g., [5]). We propose an efficient way of calculating the factor 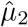 so that the entire LD matrix need not be computed.

From a theoretical point of view, following [6], we show that the GWASH estimator is consistent as *m* and *n* increase to a limiting fixed ratio, which could be greater than 1. We also provide a formula for estimating the asymptotic standard error. Simulations are used to show the statistical and computational performance of the estimator in non-asymptotic settings.

As an alternative method, Linkage Disequilibrium Score (LDSC) regression [2] has become the most popular approach for estimating heritability from summary statistics. LDSC estimates heritability by regressing squared z-scores (per SNP univariate regression *t* or Wald statistics) on corresponding “LD Scores,” defined as estimates of the sum of squared correlations for a given SNP and all others. While an effective and computationally efficient approach, LDSC was not derived from a well-specified generative model and relies on a number of heuristics, including binning LD scores, censoring outlying values, and empirical approximations to standard errors. These features are difficult to consider analytically and limit opportunities for power analyses. To compare with LDSC, we here consider a stylized version of that estimator without binning, bootstrap and other elements that allows for an easier comparison both analytically and in small scale simulations. Our simulations show that the GWASH estimator performs similarly to the LDSC regression and is closely related analytically under some conditions, suggesting avenues to further improve the theoretical foundation for both methods.

In the rest of the paper we provide the theoretical foundation for the GWASH estimator, proving its consistency and deriving its asymptotic standard error. We confirm these properties in simulations. We illustrate the feasibility of the GWASH estimator in several real GWAS datasets and show how the asymptotic standard error formula can be used for performing power analysis in prospective GWAS.

## 2 GWAS

### 2.1 The classic polygenic model

Suppose that a continuous outcome variable or phenotype is measured together with a panel of genotype markers at *m* loci for each of *n* independent subjects. Let *y*_*i*_ and 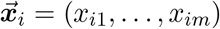 denote the outcome and genomic panel for subject *i* = 1,…,*n*. According to the classic polygenic model [9, 15], the outcome is generated according to the linear model

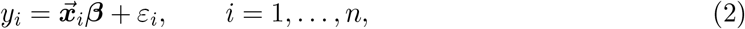

where the error terms *ɛ*_*i*_ are independent with mean 0 and variance *σ*^2^. This model may also be written in matrix form as

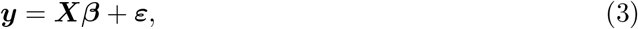

where ***y*** = (*y*1,…, *y*_*n*_)^T^, *β* = (*β*1,…, *β*_*m*_)^T^, ***ɛ*** = (*ɛ*_1_,…, *ɛ*_*n*_)^T^ and ***X*** is the regression matrix with rows 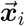, *i* = 1,…, *n*. It is also useful to write the regression matrix in terms of its columns as ***X*** = (*x*_1_,…, *x*_*m*_).

True to the sampling scheme, we shall consider the genomic panels 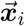 to be randomly drawn from the population together with the associated phenotypes. Let 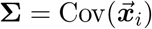 denote the *m* × *m* covariance matrix between genomic markers in the underlying population. The corresponding correlation matrix, which we shall denote 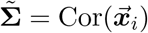, is often called the linkage disequilibrium (LD) matrix. For simplicity, our model does not explicitly include other fixed covariates (e.g. age, gender, ethnicity factors, etc.) but rather we shall assume that these other covariates have been regressed away. The interpretation of the coefficients and the heritability shall be conditional on having accounted for those other covariates and is the same as if those covariates had been included in the full model.

Similarly, rather than including an intercept term, we may equivalently assume that the vector ***y*** and the columns ***x***_1_,…, ***x***_*m*_ of ***X*** have been centered by subtracting the vector average, so that **1**^T^***y*** = 0 and **1**^T^***x***_*j*_ = 0 for *j* = 1,…,*m*. A nice consequence of centering is that, for the centered data, E(***y***) = 0 and E(***X***) = 0, where the expectation is taken with respect to the population distribution. Hence, the model (2) or (3) have no intercept term.

### 2.2 Heritability

The heritability *h*^2^ is defined as the variance explained by the predictors in model (3). Specifically, model (3) has the variance decomposition

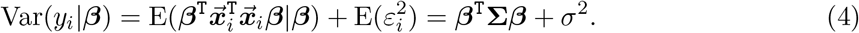

since 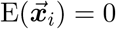. Let *τ*^2^ = *β*^T^**Σ** *β*. The heritability is the quantity [8, 15]

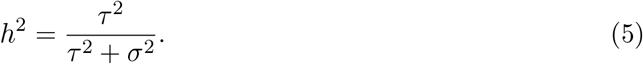

Note that in this model the vector *β* is fixed and arbitrary, with no pre-specified distribution. The model places no restrictions on the distribution of model coefficients as long as they yield the proper heritability. Thus, as opposed to other methods such as [24, 2, 25], no distributional assumptions are required on *β* in order to estimate heritability, an important point given recent debate in the literature [22].

### 2.3 GWAS univariate regressions

In GWAS, the vector of SNP effects *β* is estimated by univariate regression coefficients. Since ***y*** and the columns *x*_*j*_ are assumed centered, there is no need to fit an intercept term and the slope parameters *β*_*j*_ for each SNP *j* = 1,…,*m* are estimated via

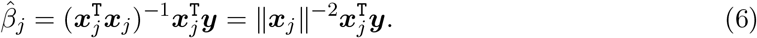

The univariate regression estimates are typically converted into t-scores by dividing by an estimate of standard error at each SNP. For each *j* = 1, …,*m*, the residual variance is

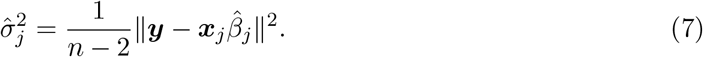

yielding the t-score

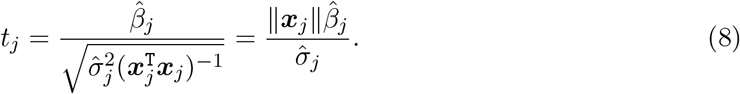

The goal is to produce an estimator of heritability that relies on the above so-called summary statistics 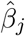, 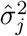 and *t*_*j*_, *j* = 1,…,*m*. We describe the heritability estimator in general in Section 3 and return to the summary statistics in Section 5.1.

## 3 The Dicker estimator

To better describe the derivation of the GWASH estimator, we first discuss the estimator proposed by [6]. Addressing the high-dimensional case where *m* is greater than *n*, [6] proposes an estimator of the fraction of variance explained by ***X*** in model (3) when the vector of coefficients *β* is fixed. While not called heritability there, this fraction is the same as the heritability defined in (5). Since ordinary least squares methods fail when *m* > *n*, Dicker’s estimator is based instead on a clever use of the method of moments. Dicker proposes two forms of the estimator depending on whether the covariance matrix **Σ**, typically unknown, is estimable or not.

### 3.1 The Dicker estimator for estimable covariance

An estimable covariance matrix **Σ** presumes the existence of a norm-consistent positive definite estimator 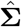, despite the dimension *m* being larger than the sample size *n*. An example of an estimable covariance matrix is a diagonal **Σ**, so that the columns of ***X*** are uncorrelated but have different variances. Other examples include matrices where the correlation structure is captured by a fixed number of parameters, such as autoregressive (AR) and exchangeable correlation models.

#### Proposition 1.

*Let* 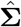 *denote the estimator of* **Σ**. *The Dicker estimator of h*^2^ *for estimable covariance [6, Sec. 4.1], denoted here 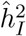, can be written (in our notation) as*

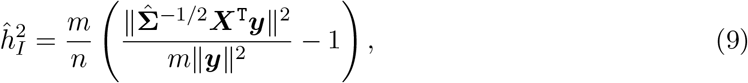

*where n is replaced by n* – 1, *owing to the centering of **y** and the columns of **X** [6, Sec. 1]*.

The estimator (9) requires a consistent estimator 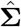 of the covariance matrix **Σ**, which is not available without further assumptions. The sample covariance matrix

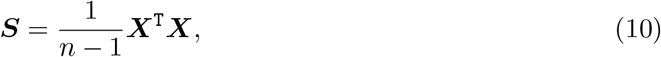

whose entries are the sample covariances of the columns of ***X***, is an unbiased estimator of **Σ**, satisfying E(***S***) = **Σ**. It is, however, not norm-consistent because the dimension *m* is larger than the sample size *n*.

Assuming that the true correlation is nonzero only close to the diagonal, as it is often assumed in GWAS, consistent estimators may be obtained, for example, by banding the sample covariance matrix [1, 3]. Even so, the estimator (9) requires computation of the inverse square root 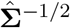. This is computationally taxing for the typical large matrix size *m* in GWAS in the order of magnitude of a million. Dicker’s estimator for unestimable covariance avoids this problem.

### 3.2 The Dicker estimator for unestimable covariance

When a model for **Σ** is not sufficiently specified to be estimable, [6] offers another form of the estimator that replaces estimation of **Σ** by estimation of its first few moments.

#### Proposition 2.

*The Dicker estimator of h*^2^ *for unestimable covariance [6, Sec. 4.2], denoted here 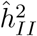, can be written (in our notation) as*

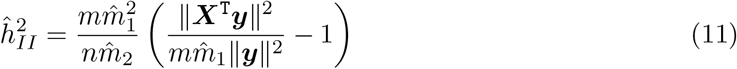

*where*

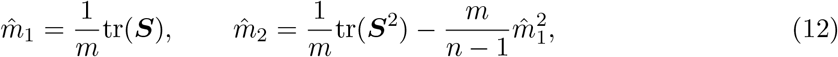

*and ***S*** is the sample covariance matrix* (10).

Proposition 2 of [6] states that if the entries of ***X*** and *ɛ* are Gaussian and **Σ** is not too far from the identity matrix (technical details omitted here), then 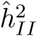 is approximately Gaussian with mean *h*^2^ and variance

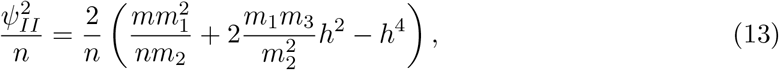

for large *m* and *n* such that *m*/*n* is bounded, where

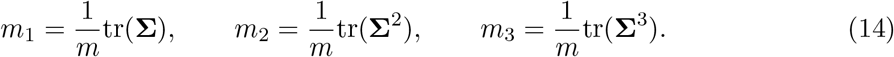

By the commutative property of the trace, it can be shown that the quantities in (14) correspond to the first, second and third moments of the eigenvalues of **Σ**. In that sense they can be called *spectral moments*.

An estimate of standard error for 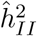 can be obtained as the square root of (13) by plugging in the estimate of *h*^2^ and those of *m*_1_ and *m*_2_ given by (12). As an estimate of *m*_3_, Dicker [6, Remark 12] suggests

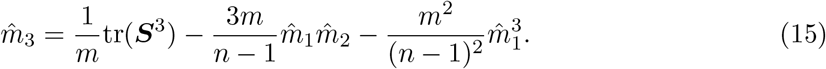

## 4 The GWASH estimator

In GWAS, it is feasible to implement Dicker’s estimator (11) if the entire dataset composed of ***X*** and ***y*** is available. However, often only GWAS summary statistics are avaiable. The GWASH estimator is essentially a modification of the Dicker estimator where the columns of ***X*** are standardized. This standardization allows writing the estimator in terms of the correlation scores defined next, which easily translate into summary statistics.

### 4.1 Correlation scores

Let 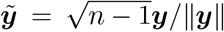 be the standardized vector ***y*** so that 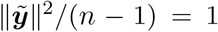. Similarly, let 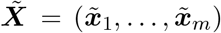 be the result of standardizing the matrix ***X*** by columns, so that the standardized columns 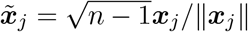 satisfy 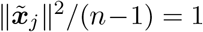, for *j* = 1,…,*m*. Because of the original centering, 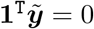 and 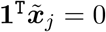.

The main idea of the GWASH estimator is to replace ***X*** and ***y*** in (11) by their standardized versions *X̃* and *ỹ*. Because (11) depends on the summary statistic ∥***X***^T^***y***∥^2^, to become 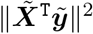, it is convenient here to define what we call the *correlation scores*

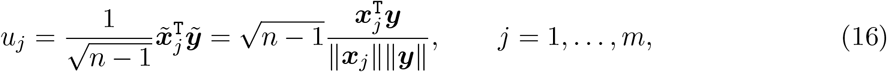

or in vector form,

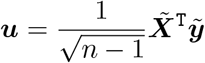

The score *u*_*j*_ may be called correlation score because it is equal to 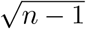 times the sample correlation between ***x***_*j*_ and ***y***. The reason for the multiplying factor is that, under the null hypothesis of no heritability (*h*^2^ = 0) so that ***x***_*j*_ and ***y*** are independent, the score (16) is asymptotically normal with mean zero and variance one. In this sense it plays the role of a z-score.

### 4.2 The LD matrix

By standardization, we may define the sample covariance matrix of the columns of *X̃*,

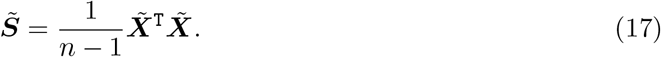

By definition, this is the sample correlation matrix with ones on the diagonal and can be referred to as the sample LD matrix.

Let 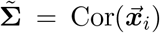 be the population correlation matrix corresponding to the covariance matrix **Σ**. Analogous to (14), we can define the first three spectral moments of Σ̃ by

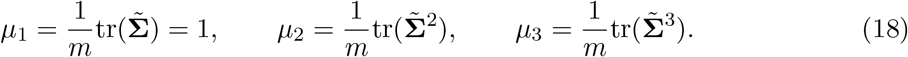

These quantities capture the total effect of LD between the genomic markers. If the genomic markers are independent with Σ̃ = ***I***, then *μ*_2_ = *μ*_3_ = 1; otherwise both moments are greater than 1.

### 4.3 The GWASH estimator from subject-level data

The GWASH estimator is defined as a modification of the Dicker estimator where: 1) ***X*** and ***y*** in (11) are replaced by their standardized versions *X̃* and *ỹ*; 2) the moment estimators 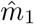, 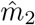 and 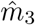 in (11), (12) and (15) are replaced by moment estimators based on the correlation matrix (17) instead. Denote the latter estimators by 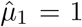, 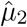 and 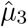; details on these are given in section 4.4 and 5.3 below.

Performing the replacements outlined above yields the expression

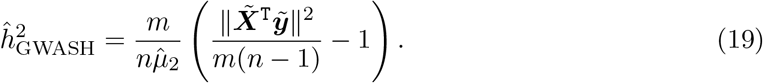

However, this expression can be written succinctly in terms of the correlation scores. We may now define our estimator.

#### Definition 1.

The GWAS heritability (GWASH) estimator is given by

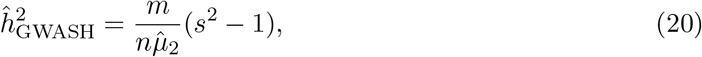

where *s*^2^ is the empirical second moment of the correlation scores:

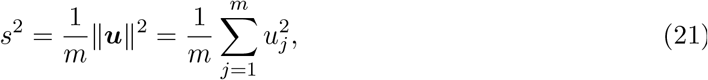

and 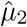 is an estimator of *μ*_2_ in (18).

The GWASH estimator depends on the data only through two summary statistics, *s*^2^ and 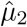. Under the null hypothesis of no heritability (*h*^2^ = 0), *s*^2^ → 1 for large *m* by the law of large numbers. Thus, (20) expresses the estimate of heritability as proportional to the excess variance of the scores with respect to the null variance 1.

The quantity 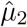 contains all the necessary information about the correlation between the predictors. From (18), if the predictors are independent, *μ*_2_ = 1. Otherwise, *μ*_2_ > 1. This implies that ignoring LD causes overestimation of the heritability. Taking account of LD is equivalent to using a smaller number of predictors. In this sense, we may define 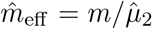 as the estimated effective number of markers and rewrite (20) as (1).

### 4.4 Estimation of the LD second spectral moment *μ*_2_

From (12), an appropriate estimate of *μ*_2_ is

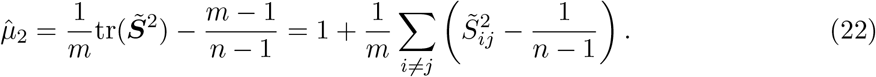

The expression on the left, in comparison to (12), uses *S̃* instead of ***S*** and uses 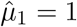 instead of 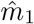. The replacement of *m* − 1 instead of *m* is more clearly understood in the expression on the right, obtained by replacing 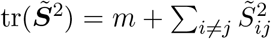. Here we can see that 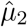 is equal to 1 (the value of *μ*_2_ under no correlation) plus 1/*m* times the total squared correlation observed in the sample LD matrix *S̃*, after subtracting from each term a bias correction of 1/(*n* − 1). The extra term 1/(*n* − 1) is the approximate second moment, for large *n*, of the empirical correlation *S̃*_*ij*_ when the true underlying correlation Σ̃_*ij*_ is zero. It is pervasive in the LD matrix and we may refer to it as a “correlation floor”.

The following lemma, whose proof is in the Appendix, states that 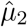 is a consistent estimator of *μ*_2_.

#### Lemma 1.

*Assume the spectral moments m*_*k*_ = tr(**Σ**^k^)/*m, k* = 1,…,4, *are bounded as m gets large. Then, as m and n get large such that m/n converges to a constant*,

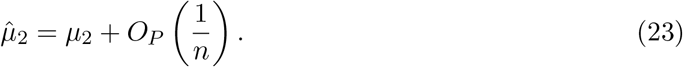

### 4.5 Asymptotic properties of the GWASH estimator

By its construction, the GWASH estimator has similar asymptotic properties to that of the Dicker estimator (11), namely consistency and asymptotic normality. Theorem 1 below shows this formally and gives the theoretical justification for using the GWASH estimator in GWAS.

#### Assumption 1.

Suppose that the assumptions of Proposition 2 of [6] hold, namely:

- The variance components *σ*^2^ and *τ*^2^, as well as the spectral moments *m*_*k*_ = tr(Σ^*k*^)/*m*, *k* = 1,…,4, are bounded.
- Let 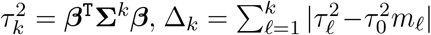, and suppose that Δ_2_ = *o*(1) and 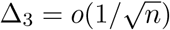.

#### Theorem 1.

*Under Assumption 1, as m and n get large such that m/n converges to a constant, the GWASH estimator* (20) *is asymptotically Gaussian with mean *h*^2^ and variance*

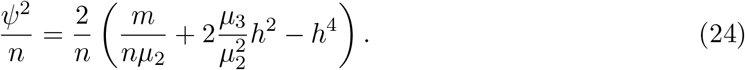

Theorem 1 implies consistency of the estimator for large *m* and *n*. The proof is given in the Appendix.

An estimate of standard error for 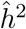 can be obtained as 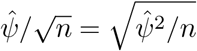, where

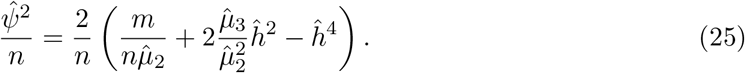

is a plug-in estimate of the asymptotic variance *ψ*^2^/*n* and 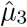 is an estimator of *μ*_3_ (see Section 5.3). The asymptotic normality of 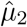 allows constructing an approximate two-sided 95% confidence interval for *h*^2^ of the form 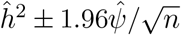. In addition, the null hypothesis *H*_0_ : *h*^2^ = 0 may be tested against the alternative *H*_*A*_ : *h*^2^ > 0 using the Wald statistic 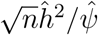 and declaring it significant at the 5% level if it exceeds the normal quantile 1.645.

## 5 Practical aspects in the context of GWAS

### 5.1 The GWASH estimator from summary statistics

In publicly available GWAS results, the original data ***y*** and ***X*** required to compute the correlation scores (16) are typically not available. Instead, it is possible to access the t-statistics (8) from the univariate regressions. Correlation scores are closely related to the t-statistics. In fact, the next result shows that the original data is not necessary, but it is possible to convert the squared t-statistics to squared correlation scores by a simple formula.

#### Proposition 3.

*The square of the correlation scores* (16) *can be obtained from the squared t-statistics* (8) *via*

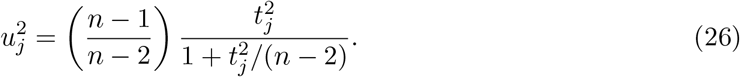

The squared correlation scores and the squared t-statistics are very close for large *n*, but not exactly. The transformation is needed because the residual variance (7) typically used in GWAS is a biased estimator of the true noise variance. The effect of the transformation is to “undo” the division by (7) and turn the t-statistic into a more appropriate score.

To compute the GWASH estimator (20), *s*^2^ can be computed directly from the u-scores (26). The LD second spectral moment 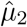 cannot be computed from summary statistics. Note, however, that 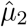 is a property of the population from which the GWAS data was sampled. Following others, we make the assumption that the sampled population has similar properties to those in public datasets such as the 1000 genomes project (1KGP) [10]. Under this assumption, 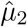 can be estimated from any random sample assayed on the same set of predictors, even if the representative sample is of a different size. For example, if a representative auxiliary dataset of size *ñ* is available on the same set of SNPs, then 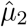 can be estimated using the methods of Section 4.4 with *ñ* instead of *n*. The same holds for 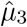 (see Section 5.3).

### 5.2 Efficient computation of the LD second moment estimator 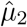

From a computational point of view, we may take advantage of the fact that, in a randomly mating population, SNPs appreciably far away within the same chromosome, or on different chromosomes, should be segregating independently. For independent markers *i,j*, their squared correlation 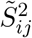 has mean of about 1/(*n* − 1), and so the terms 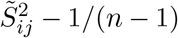 in (22) far from the diagonal are small and can be excluded from the calculation.

In general, suppose that only a set *I*_2_ of index pairs (*i,j*), *i* ≠ *j*, are included in the calculation of 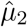. This results in the modified estimator

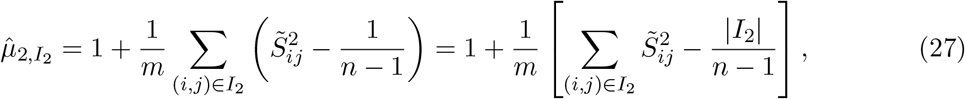

where |*I*_2_| is the number of elements in the set *I*_2_. Note that the bias correction of 1/(*n* − 1) is only applied to each of the terms actually included in the sum.

Specifically, for a single chromosome with *m*_*k*_ markers, *k* = 1,…,*K*, excluding all pairs more than *q* > 0 indexes away is equivalent to applying formula (22) to the restricted matrix 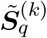 with entries

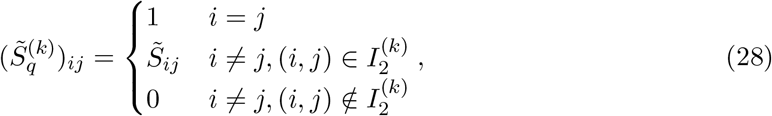

where 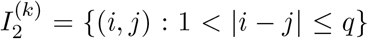. It can be shown that 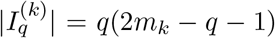, yielding the formula

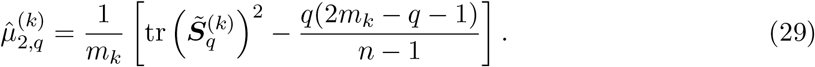

In practice, the restricted matrix (28) can be stored as a sparse matrix and the trace above computed using the property that for any squared matrix ***A***, 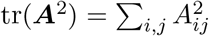.

For any set of *K* chromosomes with *m*_1_ +…+ *m*_*K*_ = *m*, it can be shown using (27) that the overall estimate 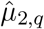 is the weighted average of the per-chromosome estimates (29), weighted by the number of markers *m*_*k*_ in each chromosome. In what follows, we refer to the distance *q* as *correlation bandwidth*.

### 5.3 Estimation of the LD third spectral moment *μ*_3_

To compute the variance (25), we need an estimator of *μ*_3_. From (15), an appropriate estimate of *μ*_3_ is

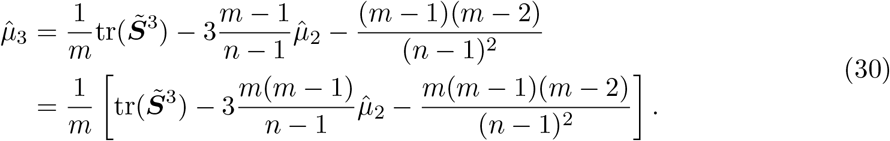

To understand this estimator, we realize that

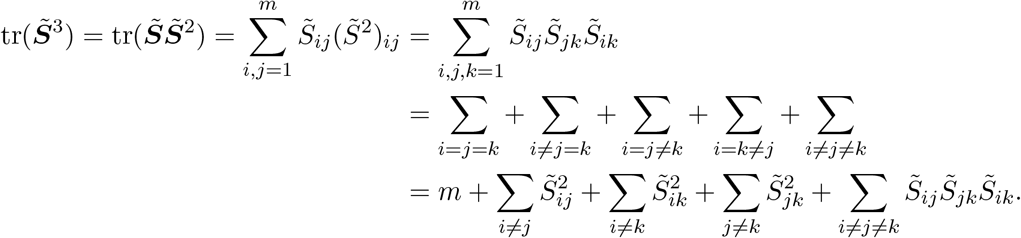

Thus, the first subtracted term in (30) makes a bias correction of 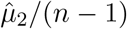 for each of the 3*m*(*m* − 1) second order terms in the sum above, while the second subtracted term makes a bias correction of 1/(*n* − 1)^2^ for each of the *m*(*m* − 1)(*m* − 1) third order terms.

As before, an approximation can be obtained by only considering entries of *S̃* close to the diagonal. Let *I*_2_ and *I*_3_ be respectively the set of index pairs (*i,j*), *i* ≠ *j*, and index triplets (*i, j, k*) other than *i* = *j* = *k* to be included in this calculation. Then we have the modified estimator

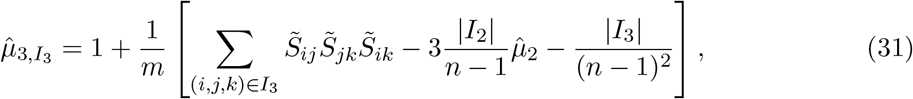

where |*I*_2_| and |*I*_3_| are the number of elements in the sets *I*_2_ and *I*_3_, respectively.

Specifically, for a single chromosome with *m*_*k*_ markers, let *S̃*_*q*_ be the restricted matrix (28) so that

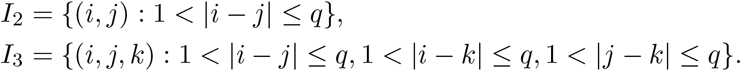

the number of elements |*I*_2_| and |*I*_3_| can be computed exactly as

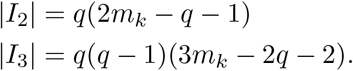

Replacing in (31) gives the formula

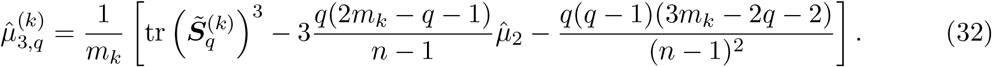

Note that the trace above can be computed using the property that for any squared matrix ***A***, tr(***A***^3^) = Σ_*i,j*_ *A*_*ij*_ (***A***^2^)_*ij*_.

Again, for *K* chromosomes, the overall estimate 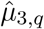 is the weighted average of the per-chromosome estimates (29), weighted by the number of markers *m*_*k*_ in each chromosome.

### 5.4 Relationship to LDSC regression

The LDSC regression method [2] is derived under different modeling assumptions to ours, the most important being the assumption that the *β* coefficients are random. In addition, the corresponding software is written for large GWAS applications and is not amenable to smaller scale simulations as we do here. To allow direct comparison both analytically and in simulations, here we consider a stylized version of LDSC regression that closely matches the GWASH estimator.

In its most basic version, written in our notation and based on our best understanding of [2], the LDSC method is essentially based on the approximation

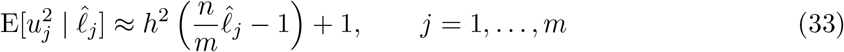

where 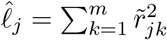 are the so-called LD-scores and *r̃*_*jk*_ are the entries of *S̃* = *X̃*^T^*X̃*/(*n*−1) (see Appendix A.4 for details). The LDSC method estimates *h*^2^ by fitting a linear model based on (33) plus observation noise. Defining 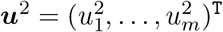 and 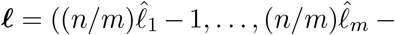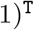, the model (33) reads E(***u***^2^) = *h*^2^*ℓ* + 1, leading to the least squares estimator

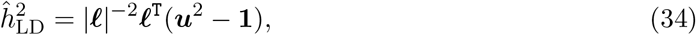

fitted with a fixed intercept equal to 1.

The GWASH estimator is related to the LDSC regression estimator above in the following way. In linear regression, the fitted line always goes through the average of the point cloud. Therefore, the average 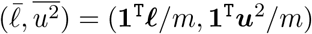 must satisfy the equation

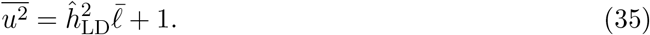

But 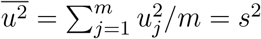 by (21) and

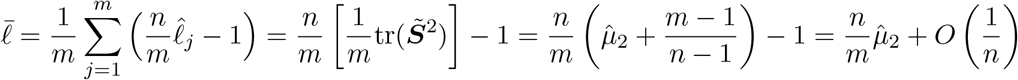

by (22), assuming that *m*/*n* converges to a constant. Replacing in (35) and comparing to (20), we obtain that

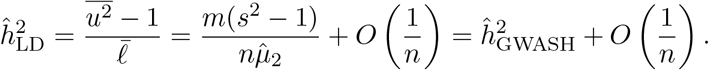

In other words, the LDSC estimator with fixed intercept equal to 1 and the GWASH estimator are asymptotically equivalent.

Another way of understanding the above derivation is that, if we consider a scatterplot of 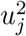 as a function of the LD scores 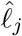, LDSC fits the least-squares straight line through the distribution, while GWASH targets the mean of the distribution directly. We will see in the simulations and data analysis that both give similar estimates but have different standard errors.

## 6 Finite sample performance

The purpose of the following simulations is to evaluate the statistical and computational performance of the GWASH estimator under various finite sample scenarios. To easily control the amount of correlation off the main diagonal, we use an autoregressive (AR) structure for the correlation matrix *S̃*.

### 6.1 Estimation of heritability

To push the limits of the estimator, in this section we consider an AR covariance structure for the predictor matrix ***X*** where the AR parameter ranges from 0 to 0.8 and where the variances of the columns of ***X*** have a wide spread from 1 to *m*. We consider two different distributions for the entries of ***X***:

- The rows of ***X*** are i.i.d. multivariate normal with mean 0 and covariance matrix **Σ** = [Diag(**Σ**)]^1/2^**Σ̃**(*ρ*)[Diag(**Σ**)]^1/2^, where Diag(**Σ**) = Diag(1,2,…,*m*) and **Σ̃**(*ρ*) is the *m* × *m* AR correlation matrix with entries Σ̃ = *ρ*^|*i*−*j*|^.
- The rows of ***X*** are i.i.d. multivariate binomial, generated using a Gaussian multivariate copula [11, 12]. According to this method, a multivariate normal vector is generated with the same covariance matrix **Σ** as above and AR parameter denoted *ρ*\*. The multivariate normal vector is then transformed to binomial by a quantile transformation with the corresponding variance. Because of the copula, the correlation matrix that is obtained for the binomial variables is not exactly AR with the specified parameter. As a reminder, we use the notation *ρ*\* to indicate the AR parameter used as an input to the multivariate copula generating mechanism.

To simulate possible structures of the vector of coefficients *β*, we consider two different scenarios for this vector:

- *β* is a single realization of *m* i.i.d. *N*(0,1) variables.
- *β* is a mixture, containing 90% of 0’s and 10% i.i.d. *N*(0,1) variables.

In all cases, the outcome ***y*** is generated according to model (3) with i.i.d. Gaussian errors. Given *β* and **Σ**, for any desired heritability *h*^2^ > 0, the error variance is set to *σ*^2^ = *τ*^2^(1 − *h*^2^)/*h*^2^ so that (5) gives heritability *h*^2^. For *h*^2^ = 0, we set *β* = 0.

Figures 1–3 show the estimates of *h*^2^ under the aforementioned combinations. The estimation methods shown are:

- The GWASH estimator (20) using the full sample correlation matrix (*q* = *m* − 1) to estimate *μ*_2_, as in (22).
- The GWASH estimator (20) using only *q* off-diagonals of the sample correlation matrix to estimate *μ*_2_, as in (29) (considered only when *ρ* > 0).
- The Dicker estimator for unestimable covariance (11).
- The simple LD regression estimator (34).

All *h*^2^ estimates are hardly distinguishable and close to the true values (grey diagonal line) within simulation error. This is precisely the desired behavior, as it shows that the GWASH estimator can estimate heritability just as well as the Dicker and LDSC estimators using only summary statistics. Note too that the correlation bandwidth *q* has little influence on the results.

### 6.2 Estimation of spectral moments and standard error

**Table 1.** Estimates of *μ*_2_ and *μ*_3_ (*m* = 1000): values presented are the mean and empirical standard deviation over 100 repetitions. The symbol *ρ*\* represents the AR parameter of the Gaussian copula.

| $\mathbf{X}$ | AR | $\mu_2$ | $\hat{\mu}_2$ | $\hat{\mu}_{2,q}$ | $\mu_3$ | $\hat{\mu}_3$ | $\hat{\mu}_{3,q}$ |
| --- | --- | --- | --- | --- | --- | --- | --- |
| | | true | Full $\tilde{\mathbf{S}}$ | Partial $\tilde{\mathbf{S}}$ | true | Full $\tilde{\mathbf{S}}$ | Partial $\tilde{\mathbf{S}}$ |
| normal | $\rho = 0.8$ | 4.55 | 4.53 (0.03) | 4.50 (0.02)<sub>$q=11$</sub> | 30.49 | 30.2 (0.48) | 29.1 (0.32)<sub>$q=11$</sub> |
| | $\rho = 0.4$ | 1.38 | 1.38 (0.006) | 1.38 (0.003)<sub>$q=5$</sub> | 2.36 | 2.35 (0.04) | 2.35 (0.01)<sub>$q=5$</sub> |
| | $\rho = 0.2$ | 1.08 | 1.08 (0.004) | 1.08 (0.001)<sub>$q=3$</sub> | 1.26 | 1.26 (0.02) | 1.26 (0.004)<sub>$q=3$</sub> |
| | $\rho = 0$ | 1 | 1.00 (0.004) | | 1 | 1.00 (0.01) | |
| binomial | $\rho^* = 0.8$ | 2.54 | 2.55 (0.02) | 2.55 (0.02)<sub>$q=11$</sub> | 10.42 | 10.48 (0.24) | 10.28 (0.18)<sub>$q=11$</sub> |
| | $\rho^* = 0.4$ | 1.13 | 1.13 (0.005) | 1.13 (0.003)<sub>$q=5$</sub> | 1.44 | 1.44 (0.02) | 1.44 (0.009)<sub>$q=5$</sub> |
| | $\rho^* = 0.2$ | 1.03 | 1.03 (0.004) | 1.03 (0.001)<sub>$q=3$</sub> | 1.08 | 1.08 (0.01) | 1.08 (0.003)<sub>$q=3$</sub> |
| | $\rho^* = 0$ | 1 | 1.00 (0.004) | | 1 | 1.00 (0.01) | |

To understand the effect of LD on the spectral moment estimators, estimates of *μ*_2_ and *μ*_3_ are shown in Table 1 under the different ***X*** structures considered above. Both 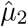 and 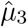 match their true values whether the full or partial sample correlation matrix is used in their estimation. Note that the empirical SE when using the partial ***S̃*** is slightly smaller than that when using the full ***S̃***.

Finally, Figures 4-6 compare estimates of standard error (SE) according to the following methods:

- Empirical SE from 100 repetitions of the GWASH estimator (20) using the full sample correlation matrix (*q* = *m* − 1) to estimate *μ*_2_.
- Empirical SE from 100 repetitions of the GWASH estimator (20) using only *q* off-diagonals of the sample correlation matrix to estimate *μ*_2_ (considered only when *ρ* > 0).
- The theoretical asymptotic SE of the GWASH estimator (square root of (25)), using the full sample correlation matrix (*q* = *m* − 1) to estimate *μ*_2_ and *μ*_3_.
- The theoretical asymptotic SE of the GWASH estimator (square root of (25)), using only *q* off-diagonals of the sample correlation matrix to estimate *μ*_2_ and *μ*_3_.
- Empirical SE from 100 repetitions of the simple LDSC regression estimator (34).
- Theoretical SE of the simple LD regression estimator (34) obtained from the linear model fit.

**Figure 1:**
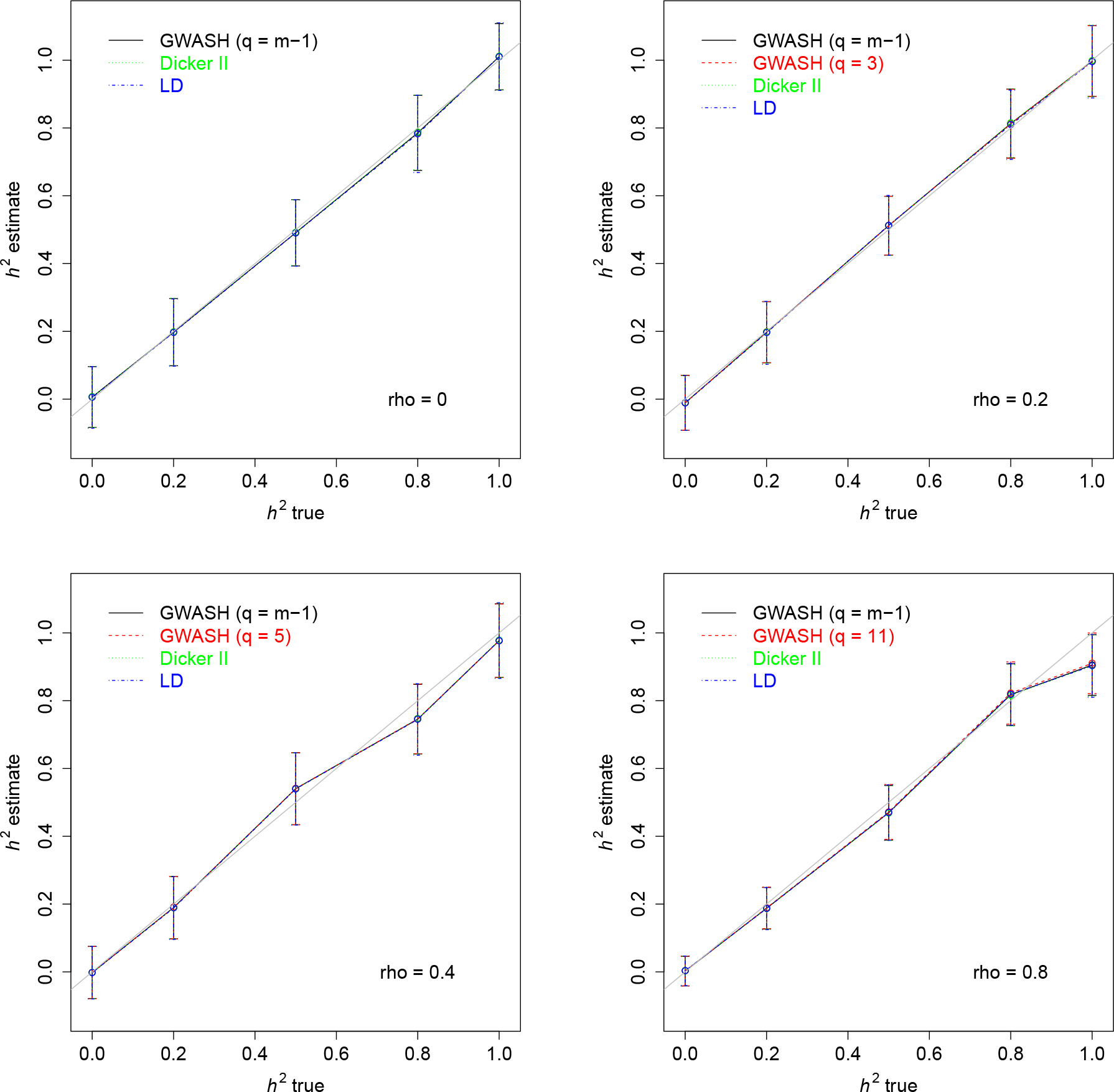
Estimates of *h*^2^ for *β* normal and ***X*** normal (*m* = 1000, *n* = 500, 100 repetitions). The bars represent the empirical standard deviation of the estimates over 100 repetitions.

**Figure 2:**
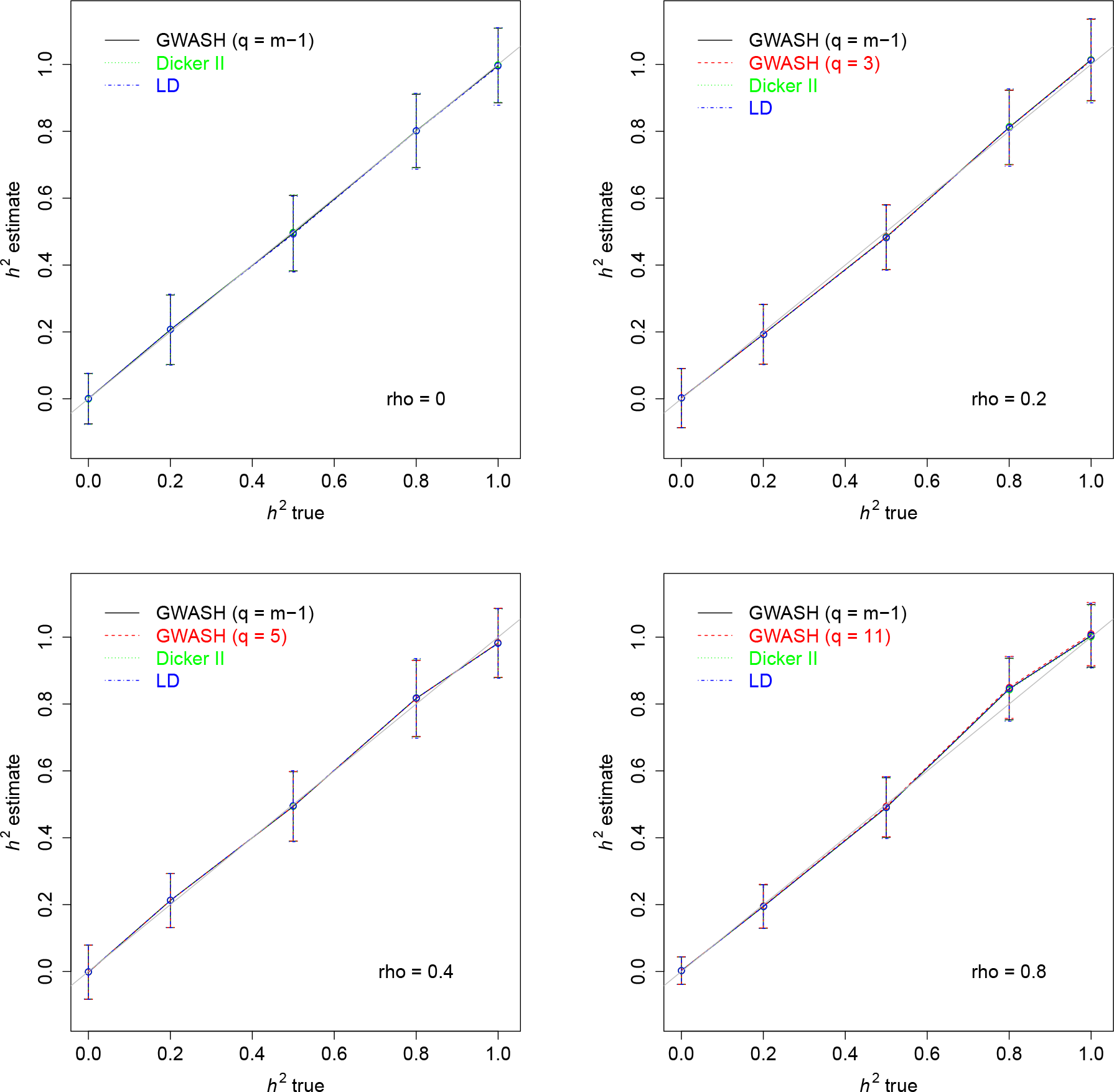
Estimates of *h*^2^ for *β* mixture and ***X*** normal (*m* = 1000, *n* = 500, 100 repetitions).

**Figure 3:**
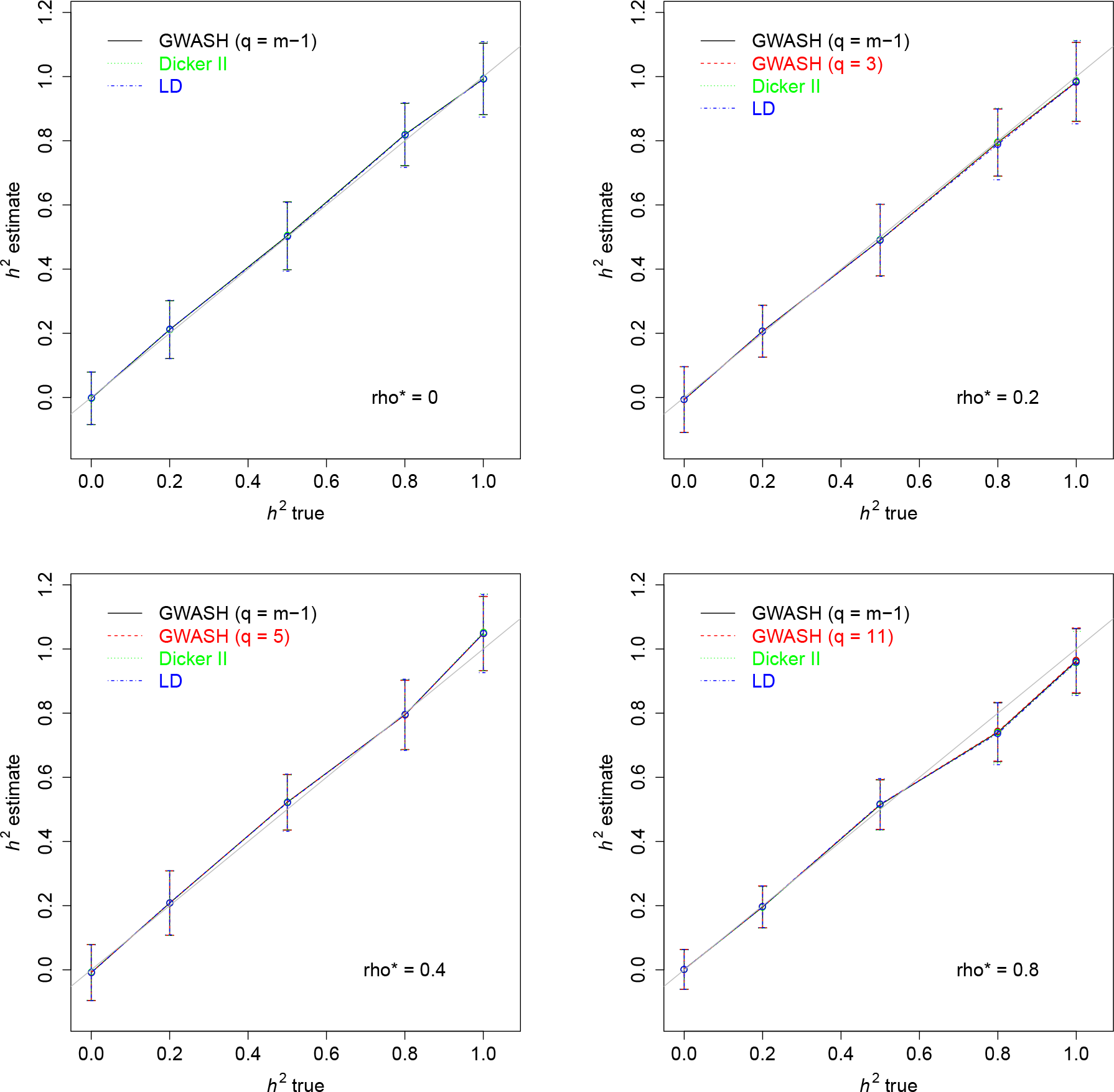
Estimates of *h*^2^ for *β* mixture and ***X*** binomial (*m* = 1000, *n* = 500, 100 repetitions).

**Figure 4:**
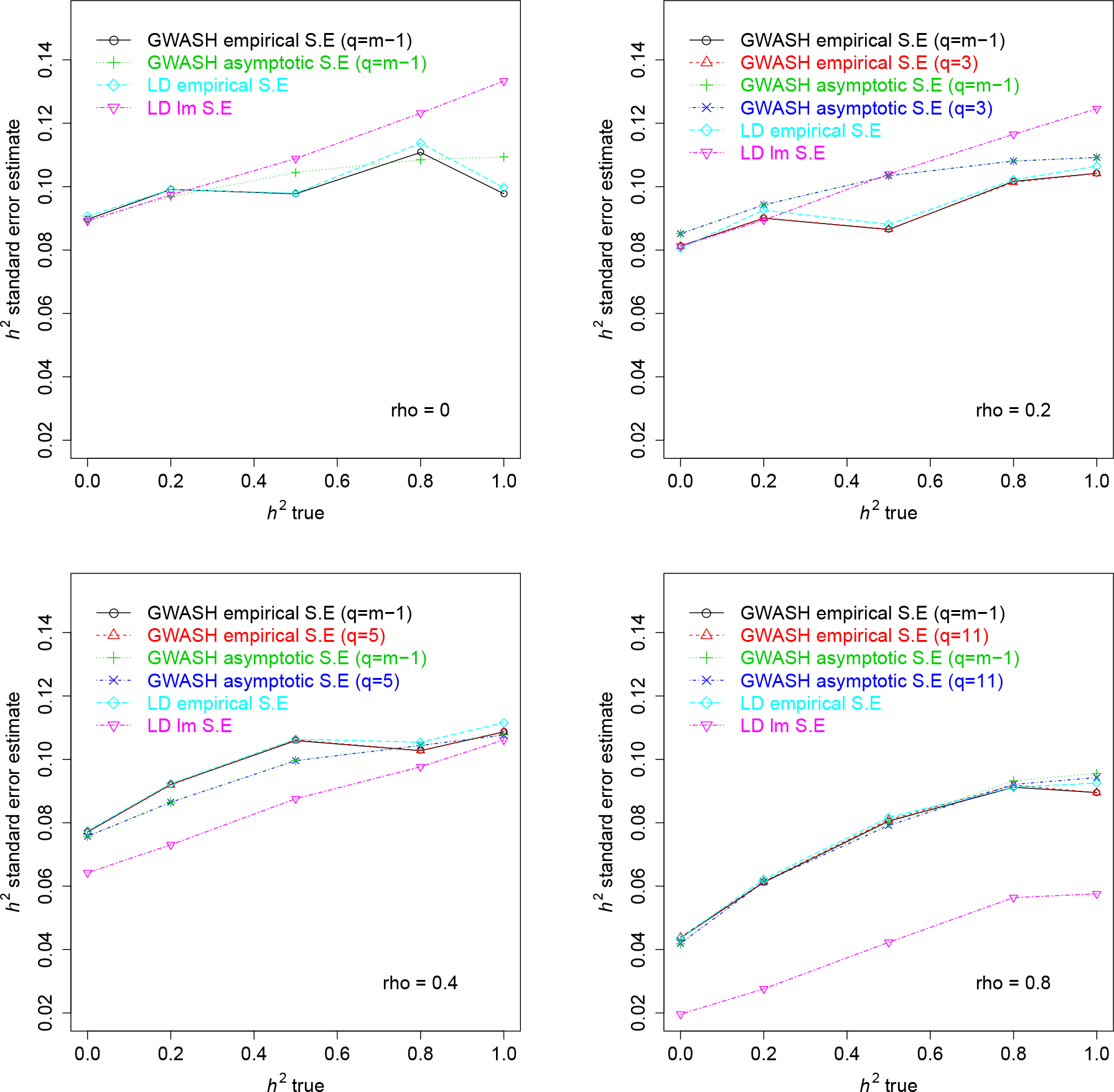
Standard errors of *h*^2^ estimates for *β* normal and ***X*** normal (*m* = 1000, *n* = 500, 100 repetitions).

**Figure 5:**
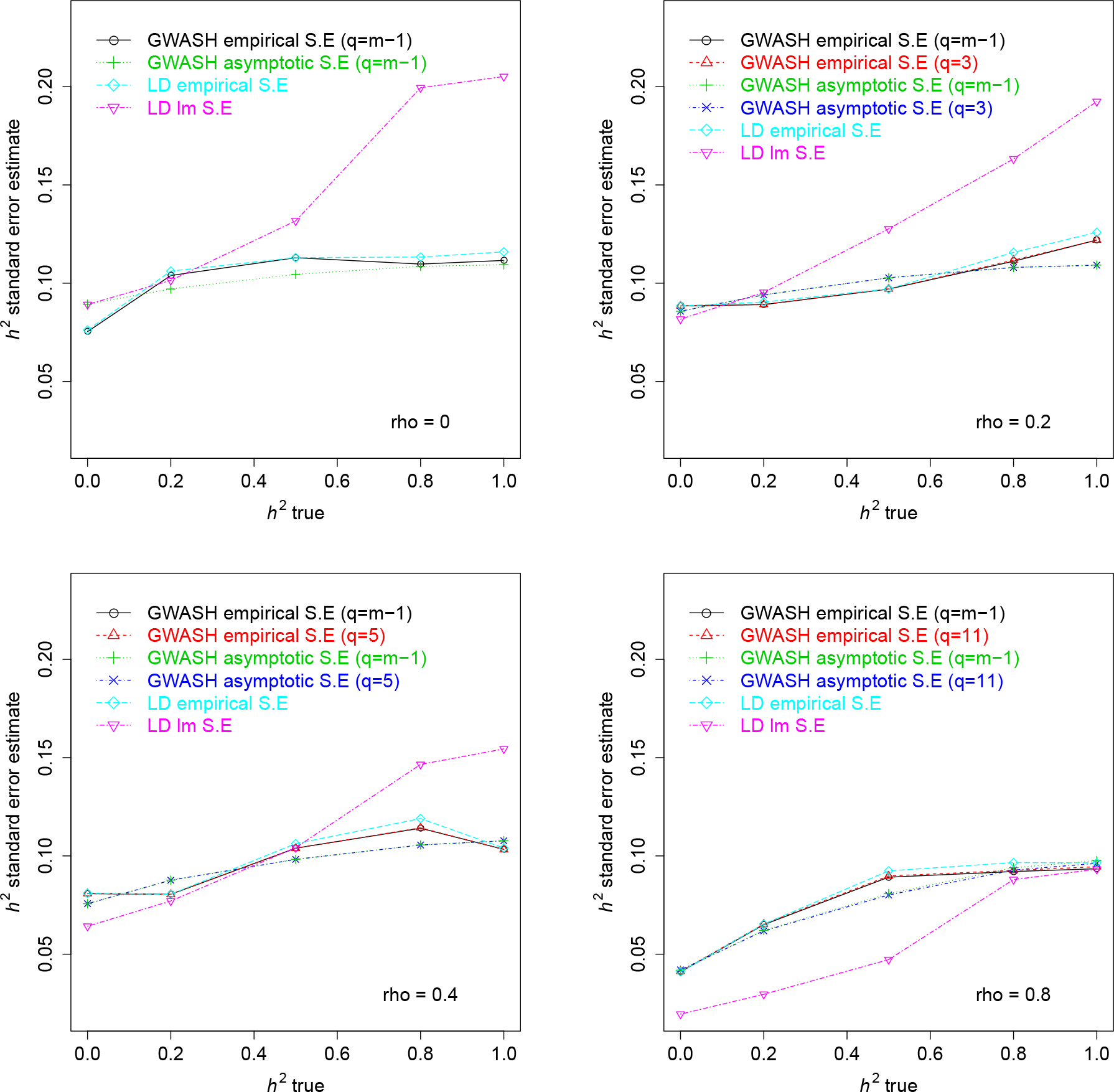
Standard errors of *h*^2^ estimates for *β* mixture and ***X*** normal (*m* = 1000, *n* = 500, 100 repetitions).

**Figure 6:**
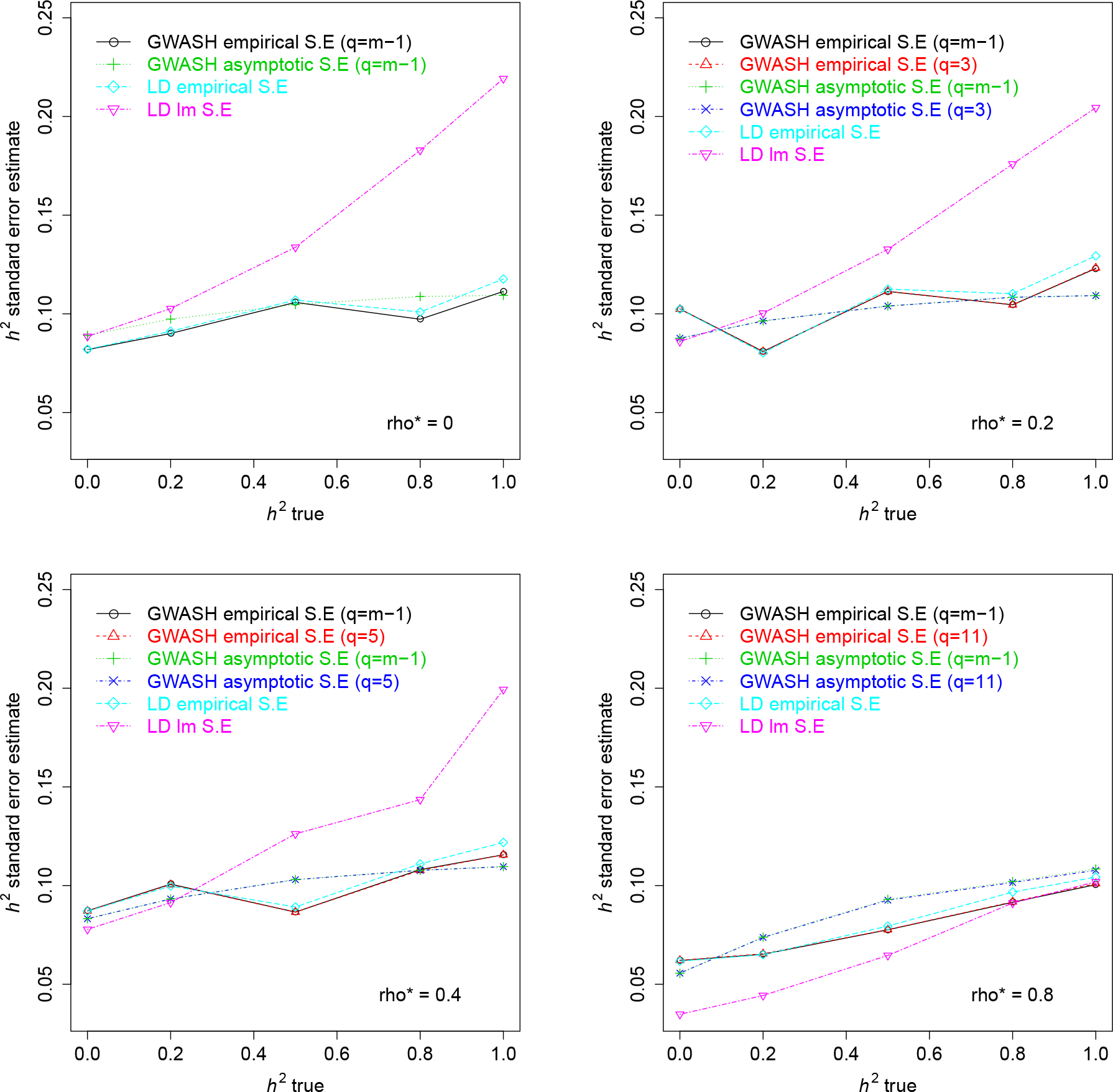
Standard errors of *h*^2^ estimates for *β* mixture and ***X*** binomial (*m* = 1000, *n* = 500, 100 repetitions).

**Table 2.**
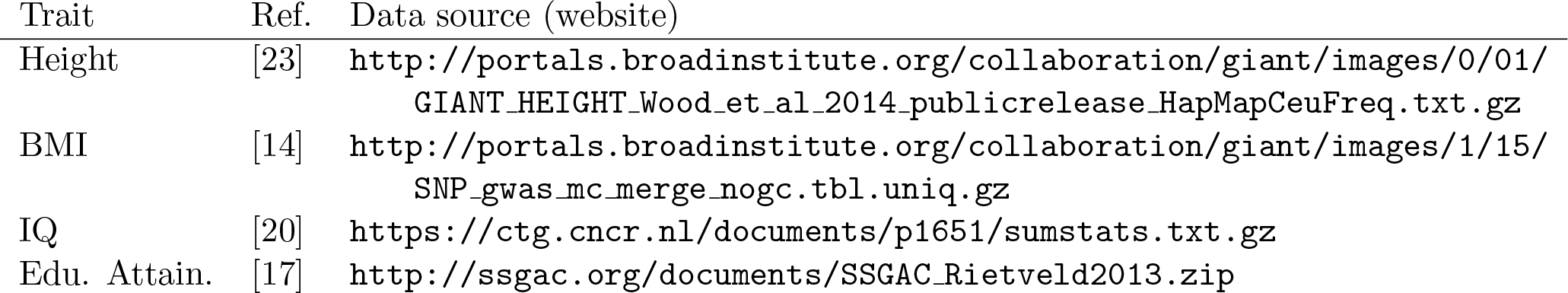
Reference information on four different GWAS studies.

In all plots, the asymptotic SE formula for the GWASH estimator approximates the empirical SE closely. For LD regression, however, overestimates or underestimates the corresponding empirical SE, explaining why the estimation of SE in [2] requires computational methods such as jackknife and bootstrapping to estimate the SE more accurately.

## 7 Real data examples

To demonstrate the performance of our estimator on real GWAS data, we estimated *h*^2^ from publicly available GWAS summary statistics for four phenotypes, height [23], body mass index (BMI) [14], IQ [20] and educational attainment [17] (Edu. Attain.). The results of the GWASH estimator were compared to the LDSC estimator [2] with intercept = 1, as follows.

### 7.1 Pre-processing of GWAS summary statistics

Summary statistics from the four GWAS studies listed in Table 2 were downloaded from the authors cited public repositories. For each study we kept the SNP name, effect allele (A1), non-effect allele (A2), per SNP sample size (*n*), association p-value (*p*) and corresponding test statistic (*t*). Where per SNP sample sizes were not available (Edu. Attain.), we used the sample size reported in the paper for each SNP. Where test statistics were not reported (BMI, Edu. Attain, Height), we converted two-tailed *p*-values to z-scores via the inverse of the normal CDF, maintaining the sign from the regression coefficients.

**Table 3:**
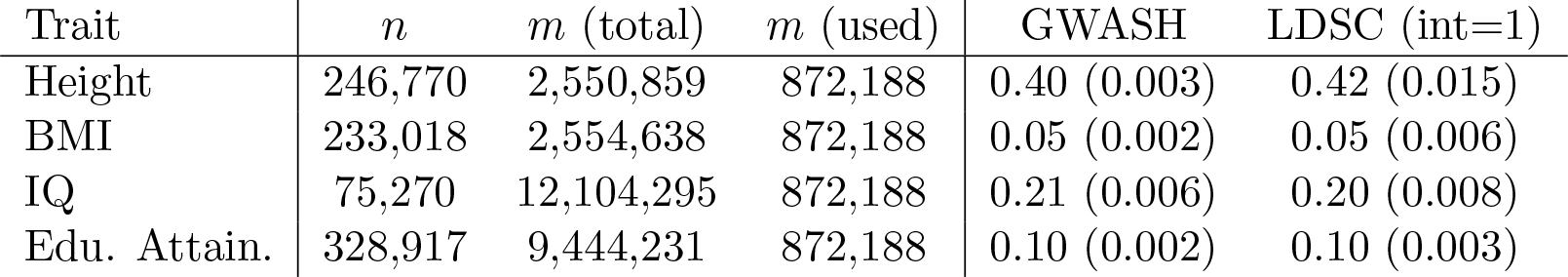
Heritability estimates for the traits in Table 2. Estimated SEs are given in parentheses.

To enable comparisons between our approach and LDSC, we kept only the subset of SNPs that was present in each of the four GWAS, had an LDSC authors precomputed LD score, and had genotype data available in the 1KGP data. We also excluded SNPs with a minor allele frequency less than 0.1% in any of the five 1KGP European subpopulations as these may be less reliably genotyped or more population variable, limiting their representativeness. This left a common set of 872,188 SNPs for each GWAS study that could be analyzed with each approach.

### 7.2 GWASH estimates and comparison to LDSC

To perform LDSC regression for each of the four GWAS studies, we downloaded all necessary software and reference data from the authors repository (https://github.com/bulik/ldsc). Both GWASH and LDSC require information about the LD among SNPs that is not typically made available alongside summary statistics. The LDSC authors address this by providing precomputed LD scores estimated from a subset of representative individuals genotypes available as part of the 1KGP data, along with their software (https://github.com/bulik/ldsc). We used these pre-computed values and their recommended protocols as faithfully as possible, following their tutorial provided (https://github.com/bulik/ldsc/wiki/Heritability-and-Genetic-Correlation) We performed the estimation with the intercept constrained to 1, as our method has theoretical similarity to the constrained intercept case.

For our estimator, calculation of 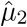 and 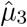 require LD information not provided with summary statistics. To compute representative values, we used a sample of the same 1KGP data (see Section 7.3 below). Table 3 shows the performance of our methods and LDSC regression on the same GWAS data. The estimated values are very similar, but the SEs for the GWASH estimator are smaller. Needless to say, all heritability estimates are highly significantly greater than zero.

### 7.3 Estimation of *μ*_2_ and *μ*_3_ from the 1KGP data

A sample of 503 individuals of European ancestry were used to compute representative values of 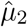 and 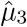. Genome-wide genotypes are available for these individuals through the 1KGP data, phase 36 (http://www.internationalgenome.org/).

The computation of 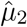 and 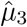 require manipulations of the LD matrix. We computed this matrix using the statistical genetic software package `plink2` [4], which provides fast routines for manipulating large genotype data sets. We can take advantage of the fact that SNPs far away from each other, or on different chromosomes are expected to have correlation (LD) approaching 0. For our reference set of genotypes we computed a banded correlation matrix with correlation bandwidth *q* = 1000 (using the notation of Section 4.4). This was done using the `plink2` commands `--r` and `--ld-window 1000`, which only returns correlations up to 1000 rows off the diagonal, for each chromosome in parallel. Matrix calculations for 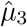 and 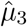 were performed in an `R` routine using sparse matrix operations in the package `matrix`. The values so obtained 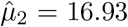 and 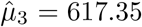 were used in the calculation of the GWASH estimate and its std. error in Table 3.

**Figure 7:**
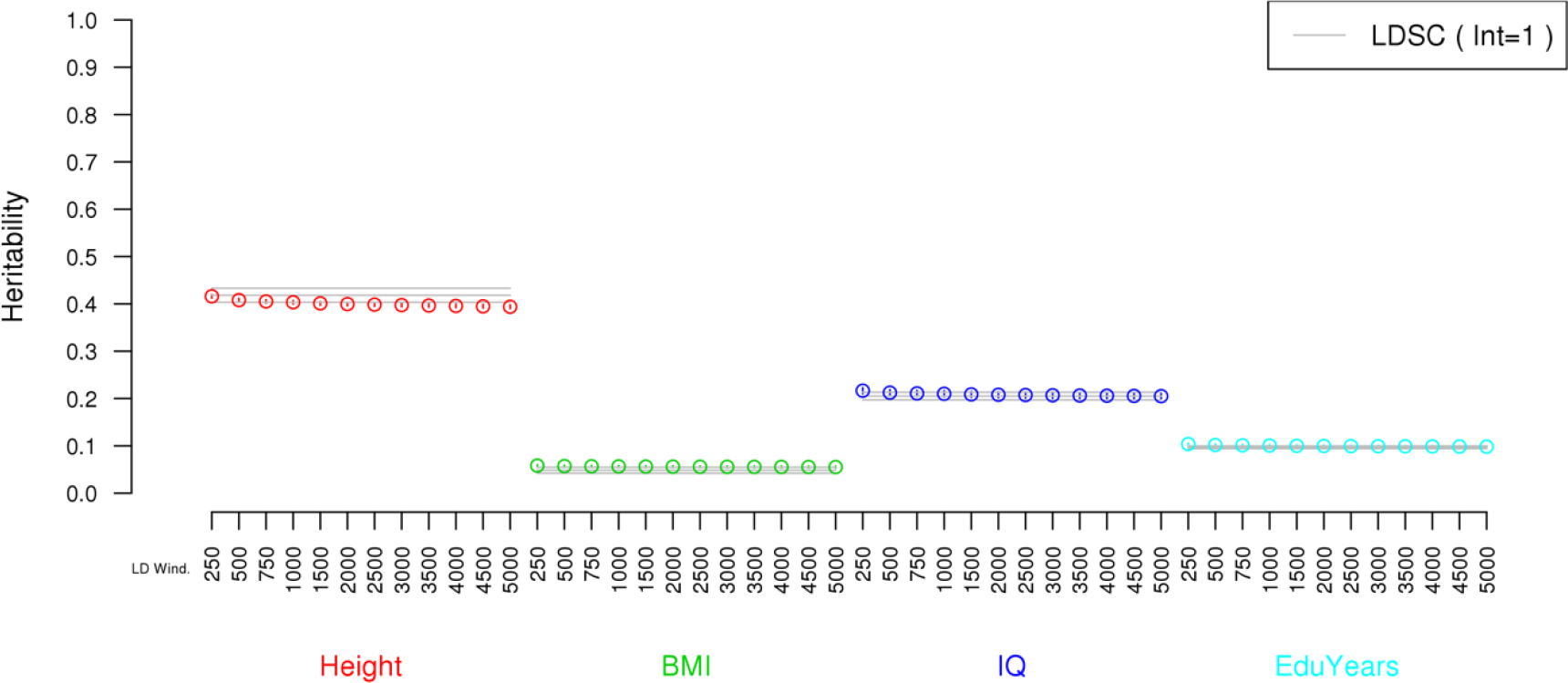
Sensitivity of GWASH estimates to the calculation of 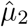 as a function of correlation bandwidth q. The LDSC (int=1) estimator is added for reference.

To evaluate the choice of correlation bandwidth *q*, the GWASH estimate was recomputed for a range of values of *q* up to 5000 used in the calculation of 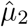. Figure 7 shows that the GWASH estimate is fairly insensitive to the correlation bandwidth *q*, the chosen value *q* = 1000 being a reasonable compromise between accuracy and computation.

## 8 Discussion

### 8.1 Estimation of SE

A nice property of the GWASH estimator, inherited from the Dicker estimator and not available with other currently used estimators, is that the precision (24) of the estimator is known theoretically based on the number of SNPs *m*, the sample size *n*, the second and third spectral moments *μ*_2_ and *μ*_3_ of the LD matrix, and the true heritability *h*^2^. The first two quantities are known from the study, while the second two can be estimated from a public resource (e.g. 1KGP). The true heritability is unknown. In this paper we have used a plug-in estimate of *h*^2^ from the study itself.

To assess the sensitivity of the SE to the value of *h*^2^, Figure 8 shows the SE (square root of (25)) as a function of the sample size *n* and the true heritability *h*^2^ using the values *m* = 872,188, *μ*_2_ = 16.93 and *μ*_3_ = 617.35 from the data analysis. The plot shows that the SE is almost insensitive to the value of *h*^2^, increasing only slightly as *h*^2^ increases for any fixed *n*. As a consequence, as an alternative to (25), a slightly conservative but more stable estimate of the SE can be obtained by plugging in the worst-case value *h*^2^ = 1 instead of the estimated value 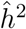.

**Figure 8:**
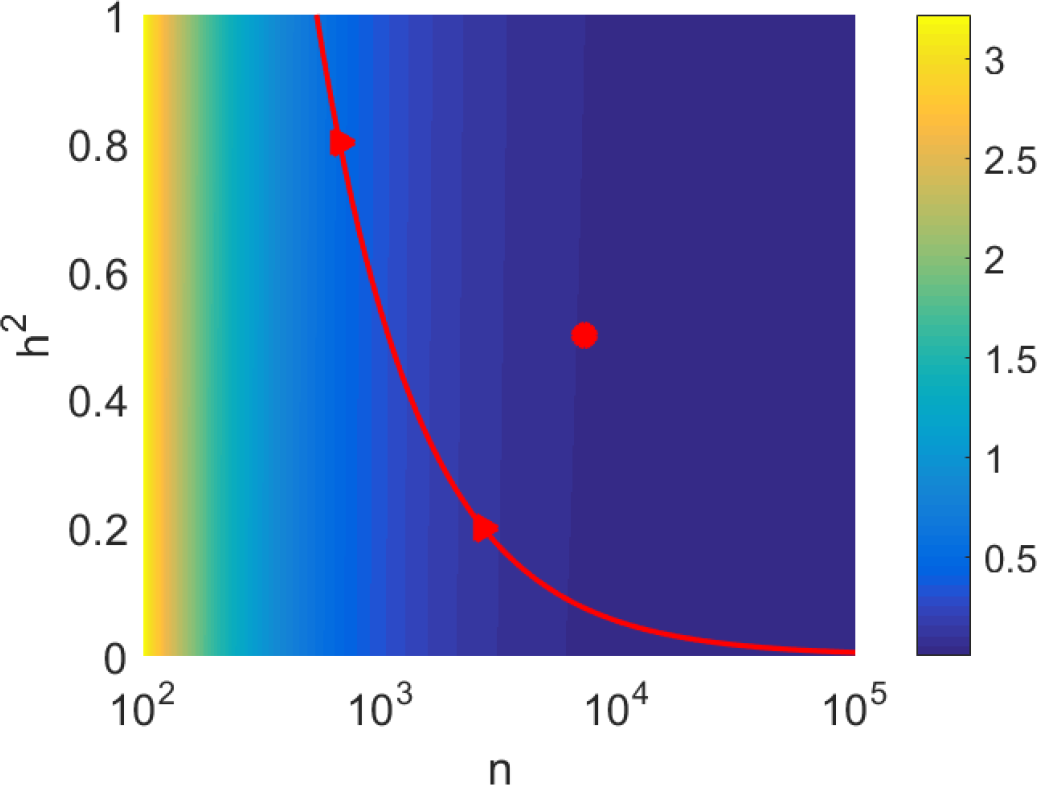
The SE of the GWASH estimator as a function of *n* and *h*^2^, for *m* = 872,188, *μ*_2_ = 16.93 and *μ*_3_ = 617.35. The red curve indicates the pairs (*n, h*^2^) for which *h*^2^ =1.645 SE; values of *n* to the right of the curve allow detection of a non-zero heritability at the 5% level.

### 8.2 Sample size and power calculations for prospective GWAS

Relation (24) can be used in a prospective study to determine the number of subjects required to estimate heritability according to a desired accuracy. Given any fixed set of *m* SNPs, the values of *μ*_2_ and *μ*_3_ may be estimated from a public resource (e.g. 1KGP) and then the SE can be designed as a function of *n* and the targeted *h*^2^. For the values of *m*, *μ*_2_ and *μ*_3_ in the data analysis, *Figure 8* shows that the SE can be quite large for small *n*, but it drops as *n* increases.

From a design point of view, the sample size *n* can be chosen to achieve a desired SE. For example, for a heritability of *h*^2^ = 0.5, an SE of 0.05 is achieved with *n* = 7234 (red circle in Figure 8). The SE can also help design studies with the goal of detecting a heritability that is significantly greater than zero. As mentioned at the end of Section 4.5, a one-sided Wald test will be significant at the 5% level if the estimate of *h*^2^ is greater than 1.645 SEs. In Figure 8, this corresponds to choosing *n* to the right of the red curve. For example, to detect a heritability of *h*^2^ = 0.8, the minimal sample size is *n* = 673; to detect a heritability of *h*^2^ = 0.2, the minimal sample size is *n* = 2699 (red triangles).

## A Proofs

### A.1 Proofs of shorter propositions

*Proof of Proposition 1.* Dicker [6, Section 4.1] proposes the estimators

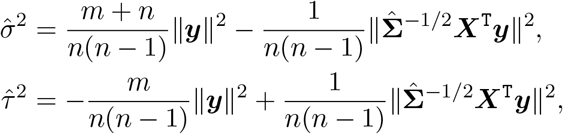

where *n* has been replaced by *n*−1 due to the centering of ***y*** and ***X***. The estimator of heritability is the fraction

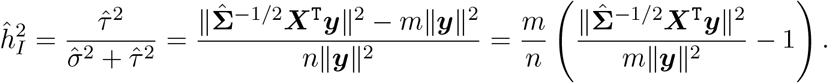

*Proof of Proposition 2.* Dicker [6, Section 4.2] proposes the estimators

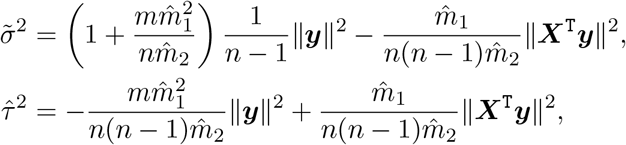

where *n* has been replaced by *n*−1 due to the centering of ***y*** and ***X***. The estimator of heritability is the fraction

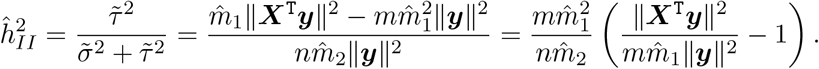

*Proof of Proposition 3.* From (7) and (8),

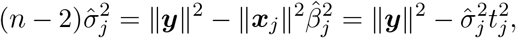

so 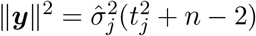. Thus, using (6) and again (8), we can write

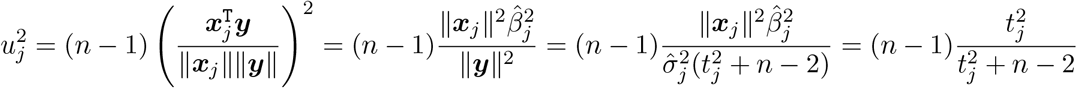

and the result follows.

### A.2 Supporting lemmas for the proof of Theorem 1

*Proof of Lemma1.* Any two sample correlations *S̃*_*jk*_ and *S̃*_*lh*_ are asymptotically bivariate normal with means Σ̃_*jk*_ and Σ̃_lh_, and variances 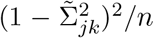 and 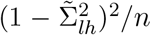. The covariance Cov(*S̃*_*jk*_, *S̃*_*lh*_), whose expression is given by [7], is a polynomial of order 4 in Σ̃_*jk*_ and Σ̃_lh_ and proportional to 1/*n*. From the asymptotic normality, we may write

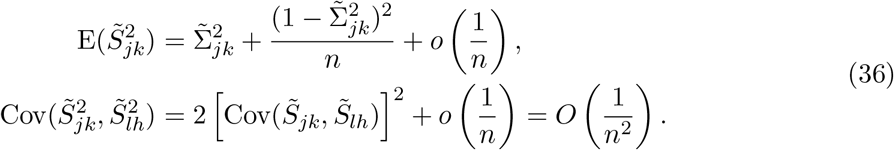

Now, from (22) and (36),

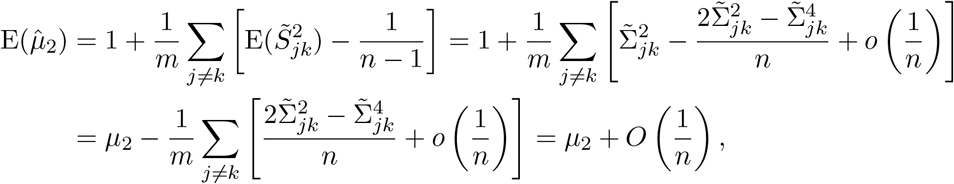

since the spectral moments of **Σ**̃ up to order 4 are assumed bounded by Assumption 1. Furthermore,

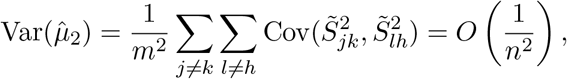

again because the spectral moments of **Σ**̃ up to order 4 are assumed bounded. Thus,

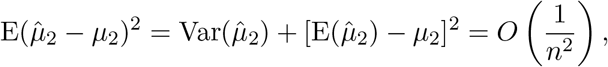

implying (23).

#### Lemma 2.

*Let **D*** = Diag(***S***) **Δ** = Diag(**Σ**). *Suppose Assumption 1 holds. Then*

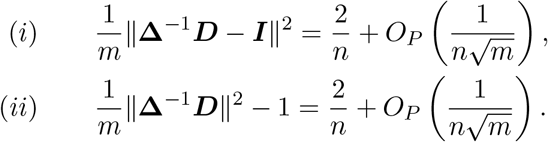

*Proof.* For the purpose of simplicity of the proof, assume that the columns of ***X*** have not been centered so that the rows of ***X*** are independent. This makes no difference asymptotically, only changing de number of degrees of freedom *n* − 1 to *n*. Let *W*_*j*_ = ∥***x***_*j*_∥^2^/[*n*Σ_*jj*_] be the *j*-th diagonal entry of **Δ**^−1^***D*** where Σ_*jj*_ is the *j*-th diagonal entry of **Σ** (or **Δ**). Then 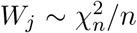 has mean 1 and variance 2/*n*.

i. The expectation of 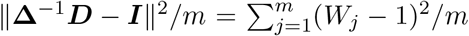 is equal to Var(*W*_*j*_) = 2/*n*. Its variance is

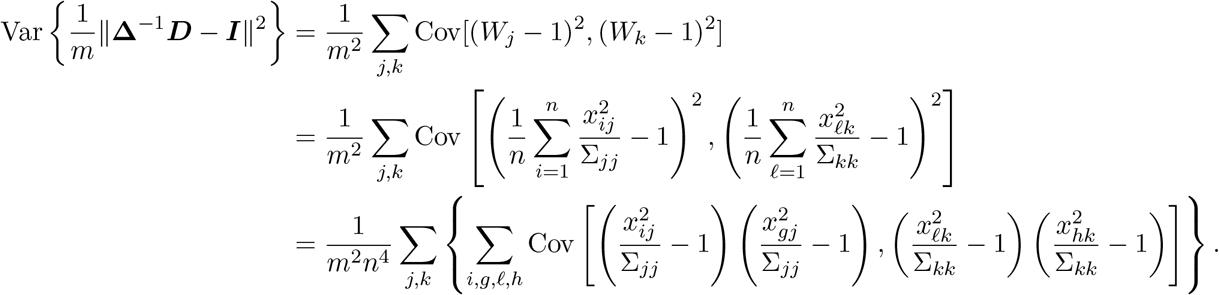

Since the rows of ***X*** are independent, most terms in the inner sum vanish except for the cases *i* = *g* = *ℓ* = *h*, where the inner covariance is 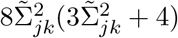 (and Σ̃_*jk*_ is the (*j, k*) entry of **Σ**̃), and the terms *i* = *ℓ* ≠ *g* = *h* and *i* = *h* ≠ *g* = *ℓ*, where the inner covariance is 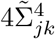. Here we have used the fact that within any row *i* of ***X***,

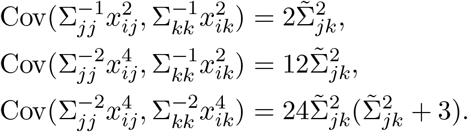

Hence,

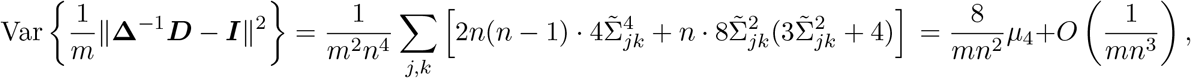

where 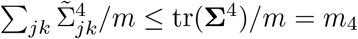 is bounded by Assumption 1. This yields (i).
ii. Write

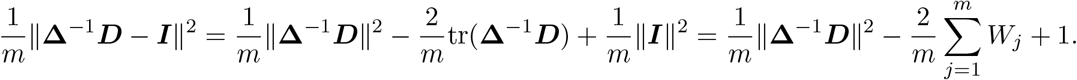

Thus ∥**Δ**^−1^ ***D***∥^2^/*m* has expectation 1 + 2/*n*. Similar to the proof of (ii), it can be shown that the variance is of the same order.

### A.3 Proof of Theorem 1

Consider a situation where, instead of ***X***, the observed covariate matrix is ***Z*** = ***X*** **Δ**^−1/2^ with **Δ** = Diag(**Σ**). The idea of the proof is to approximate the GWASH estimator (20) by an estimator based on ***Z***. This is done in two steps by: (1) establishing the asymptotic properties of the estimator based on ***Z***; (2) establishing the asymptotic equivalence of the two estimators.

Let ∥*A*∥^2^ = tr(*A*^T^*A*) denote the squared Frobenius norm of the matrix *A* and recall the Cauchy-Schwartz inequality tr(*A*^T^*B*) ≤ ∥*A*∥∥*B*∥.

<u>Step 1:</u> Based on ***Z***, the full model (3) can be written as

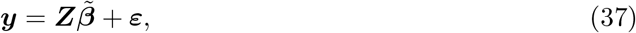

where *β*̃ = **Δ**^−1/2^***β***. Note that *τ*^2^ = ***β***^T^**Σ***β* = *β*̃^T^**Σ**̃*β*̃, so the heritability (5) does not change. The matrix ***Z*** has rows with covariance **Σ**̃ = **Δ**^−1/2^**Σ Δ**^−1/2^ and its sample covariance matrix is ***Z***^T^***Z***/(*n* − 1) = **Δ**^−1/2^***S*****Δ**^−1/2^. Following the form of Dicker’s estimator (11) for unestimable covariance, the heritability estimator based on ***Z*** is

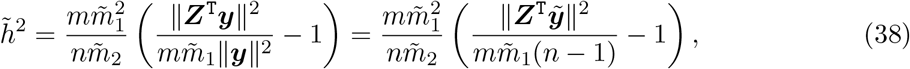

where

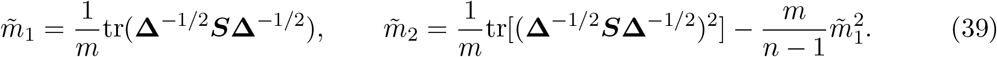

Applying Proposition 2 of [6] to the estimator (38) gives that this estimator is asymptotically Gaussian with mean *h*^2^ and variance

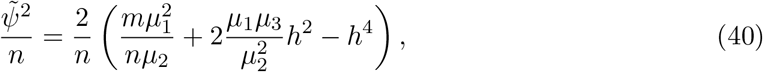

where *μ*_1_ = 1, *μ*_2_ and *μ*_3_, given by (18), are the spectral moments of the covariance of ***Z***.
<u>Step 2:</u> To show that the GWASH estimator (20) is asymptotically equivalent to the estimator (38) based on ***Z***, we need to approximate: (i) ∥***X***̃^T^***y***̃∥^2^/*m* by ∥***Z***^T^***y***̃∥^2^/*m*, and (ii) 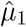 by *m*̃_1_ and 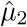 by *m*̃_2_.
  i. Let ***D*** = Diag(***S***) so that ***X***̃^*T*^ = ***D***^−1/2^***X***^*T*^ = ***D***^−1/2^**Δ**^1/2^***Z***^*T*^. Define *v* = ***Z***^T^***y***̃. We may write

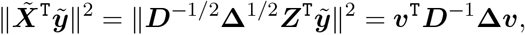

so

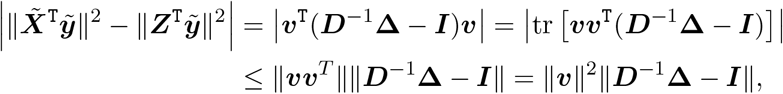

where the bars around matrices denote the Frobenius norm and the inequality is due to the Cauchy-Schwarz inequality for the Frobenius norm. By Lemma 2,

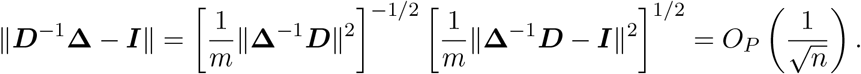

Thus

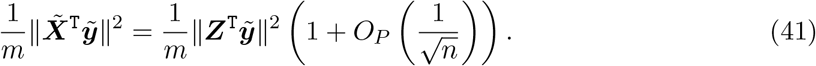
  ii. As stated in the proof of Proposition 2 of [6], applied to the estimator (38) based on ***Z***, we have that

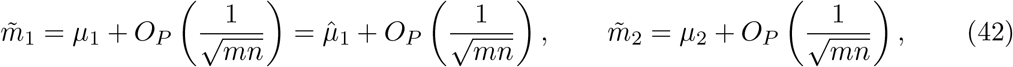

because 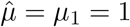. Furthermore, (23) together with (42) imply

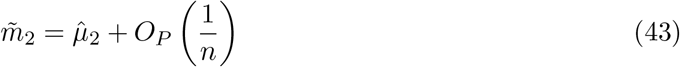

because *m*/*n* converges to a constant. Putting (41), (42) and (43) together in (38) and comparing to (19), we obtain that

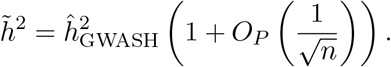

Hence 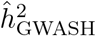 has the same asymptotic distribution as *h*̃^2^. Note that the variance (40) is the same as (24).

### A.4 Derivation of the LDSC regression equation (33)

In our notation, the chi-squared statistics defined in Section 1.2 of the Supplementary Note to [2] can be written as 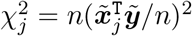, because both the predictors and the response in [2] are assumed standardized. Using (16), we can write 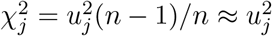 for large *n*. Then Eq. (1.3) in [2] becomes

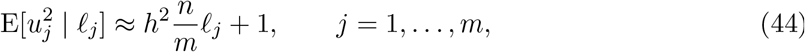

where 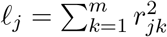 and 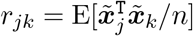 However, the Online Methods section in [2] explains that the empirical LD-scores are biased and cannot be used directly in (44). Instead, they use the adjusted LD-scores

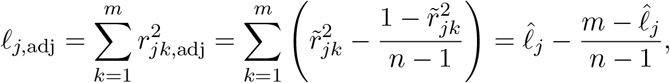

where 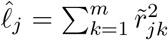 and we have used *n* − 1 in the adjustment instead of *n* − 2 to reflect the fact that the relevant LD-regression fit in our case uses a fixed intercept. Using the adjusted LD-scores in (44), we have that

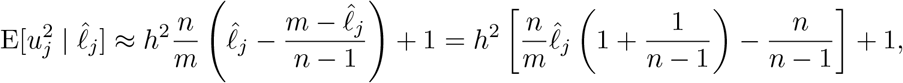

yielding (33) for large *n*.

## Acknowledgments

This work was partially supported by NIH grant 1R01GM104400 and The Lundbeck Foundation
Initiative for Integrative Psychiatric Research.

